## Supplementary information for "tRNA lysidinylation is essential for the minimal translation system found in the apicoplast of *Plasmodium falciparum*"

**Supplementary Table 1. Primers used in this study.** Restriction enzyme sites are underlined.

| **Primer name** | **Sequence (5’🡪3’)** | **Primer description** |
| --- | --- | --- |
| Primers to amplify homology arms (HA) and guide RNA (gRNA) annealing for *pftilS* knockout | | |
| TilS.HA1F | gtgccacgagcggccGCAGGTTCCATAGAAATGTTGTGTG | Forward for HA1 amplification |
| TilS.HA1R | aagcgcagcggccGCGTAAAACATCTTGATTCTTTTTAATACA | Reverse for HA1 amplification |
| TilS.HA2F | ttcgacagacgccggCAAAGGATATATGAACAGATGATAGCA | Forward for HA2 amplification |
| TilS.HA2R | tggccaccagccggCATCATGTTGTATATGTGTATCTTTTT | Reverse for HA2 amplification |
| Tils.gRNA.pF | /5Phos/tattTCAAGATGTTTTACAACTTG | 5’-phosphorylated guide RNA forward oligo |
| Tils.gRNA.pR | /5Phos/aaacCAAGTTGTAAAACATCTTGA | 5’-phosphorylated guide RNA reverse oligo |
| Primers for *pftilS* gene knockout confirmation | | |
| TilS.5.F | ATGTGTCGAAGAGATGGCAATTTTTCATACCACAT | Forward for 5’ and Δ5’ PCR |
| TilS.5.WT.R | TATGAATGAATAAAGTAGACACAAAGAATCTACCCCTGA | Reverse for 5’ PCR |
| TilS.3.WT.F | TGAACTTTTATTATTACCCTCTAAACTAATTAGACTAGAAAT | Forward for 3’ PCR |
| TilS.3.R | TACAAAAATGTTAGCACTTTTTTCTTTAAAGCACACTAATG | Reverse for 3’ and Δ3’ PCR |
| pRS.F | CATATTTATTAAATCTAGAATTCGACAGACGCCG | Forward for Δ3’ PCR |
| pRS.R | TACAAAATGCTTAAGCGCAGCGGCC | Reverse for Δ5’ PCR |
| Primers to amplify representative genes from nuclear and organellar genome | | |
| LDH.F | GGAGATGTAGTTTTGTTCGATATTG | Forward for PCR |
| LDH.R | CTTGTAAAGGGATACCACCTACAG | Reverse for PCR |
| SufB.F | CATGTAGCTATAGTAGAAATAATAGTAAAAGATTATGG | Forward for PCR |
| SufB.R | GACTCTGAAATACTTAAACCACGTTGC | Reverse for PCR |
| Cox1.F | CTTCATCTTTAAGAATAATTGCACAAGAAAATGTAAATC | Forward for PCR |
| Cox1.R | GTACATATGATGTACCCATACTAAGCTTCC | Reverse for PCR |
| Primers for generation of pCLD-*ectilS*-*mcherry-10xapt* plasmid | | |
| EcTils.InF F | GTTAGAAGGTTCCGGAATGACACTCACGCTCAATAGACAAC | Forward for *ectilS* amplification |
| EcTils.InF R | gcccttgctCgtacgactaagcgttttctgccagacaaaac | Reverse for *ectilS* amplification |
| Primers for confirmation of gene knock-in | | |
| P230p.out.HA.F | Ggttgtgatttttcaggtgattcc | Forward for attL and attB PCR |
| P230p.out.HA.R | gaaaattgtaggggcagctaaatccgac | Reverse for attR and attB PCR |
| attB.Int.F | GCAGTGTGGAATTCCCTGCA | Reverse for attL PCR |
| attB.Int.R | TTAAGTGTAGTTAATTCATCAAATAGCATGC | Forward for attR PCR |
| Primers for sequencing | | |
| pRS.R | TACAAAATGCTTAAGCGCAGCGGCC | For HA1 insertion in pRSng-*pftilS* |
| pRS.F | CATATTTATTAAATCTAGAATTCGACAGACGCCG | For HA2 insertion in pRSng-*pftilS* |
| pL6.gRNA.F | GGGTAAATTATTATTAAAAAATGTATATGTTATG | For guide RNA insertion in pCasG-LacZ |
| CLD2.F | acaacctaggATGAAGATCTTATTAC | Forward for *ectilS* insertion in pCLD-*ectilS-mcherry-10xapt* |
| RFP.R | gagggctccgtgaacggc | Reverse for *ectilS* insertion in pCLD-*ectilS-mcherry-10xapt* |
| Cre.Ins.Seq.F | CAACCtagAtaacttcgtatagcatacattatacg | Forward for *pftilS* (full-length or truncated) insertion in pCre-*tr/pftilS-3xV5* |
| TilS.Seq.F1 | GGATGTGCTGCAGTTGGTAGAAGGA | Forward for *pftilS* (full-length or truncated) insertion in pCre-*tr/pftilS-3xV5* |
| TilS.Seq.F2 | CAGTAAGGAGTACTCCAACGATTCG | Forward for *pftilS* (full-length or truncated) insertion in pCre-*tr/pftilS-3xV5* |
| TilS.Seq.F3 | GACCACGAACCTGGAAAATTACC | Forward for *pftilS* (full-length or truncated) insertion in pCre-*tr/pftilS-3xV5* |
| NewApt.5R | ctcgcTATCAAGGAATCgagtcc | Reverse for *pftilS* (full-length or truncated) insertion in pCre-*tr/pftilS-3xV5* |

**Supplementary Table 2. Proteins used for phylogenetic analysis presented in Supplementary Figure 3.** ^a^Uniprot ID, ^b^PlasmoDB ID, ^c^ToxoDB ID, ^d^PiroplasmaDB ID.

| **Species** | **Accession** | **Gene name** |
| --- | --- | --- |
| Only Syngen Nebraska virus 5 (Nv) (Chlorovirus) | A0A1J0F9X6^a^ | OS5_287L |
| *Chlorella sorokiniana* (Freshwater green alga) | A0A2P6TQ55^a^ | C2E21_4934 |
| *Gracilaria tenuistipitata* var. *liui* (Red alga) | Q6B8L1^a^ | tilS, ycf62, Grc000193 |
| *Oryza sativa* (Monocot) | B8AFB7^a^ | OsI_06667 |
| *Zea mays* (Monocot) | A0A804PGD6^a^ | tilS |
| *Zingiber officinale* (Monocot root plant) | A0A8J5GGP9^a^ | ZIOFF_041066 |
| *Arabidopsis thaliana* (Dicot) | F4J7P7^a^ | RSY3 |
| *Perkinsus olseni* (Dinoflagellate) | A0A7J6UN02^a^ | FOZ63_029171 |
| *Synechocystis* sp, PCC 6803 (Freshwater cyanobacteria) | P74192^a^ | tilS, slr1278 |
| *Vitrella brassicaformis* CCMP3155 (Dinoflagellate) | A0A0G4FYP9^a^ | Vbra_16478 |
| *Escherichia coli* (Gram-negative bacteria) | P52097^a^ | tilS, mesJ |
| *Geobacillus kaustophilus* (Thermophilic Gram-positive bacteria) | Q5L3T3^a^ | tilS, GK0060 |
| *Bacillus subtilis* (Gram-positive bacteria) | P37563^a^ | tilS, yacA |
| *Aquifex aeolicus* (chemolithoautotrophic Gram-negative bacteria) | O67728^a^ | tilS, aq_1887 |
| *Mycoplasma genitalium* (Gram-negative bacteria) | P47330^a^ | tilS, MG084 |
| *Plasmodium falciparum* (Apicomplexan) | PF3D7_0411200^b^ | tilS, PP-loop family protein |
| *Toxoplasma gondii* (Apicomplexan) | TGME49_215100^c^ | tilS, PP-loop family protein |
| *Neospora caninum* (Apicomplexan) | NCLIV_052110^c^ | tilS, hypothetical protein |
| *Eimeria tenella* (Apicomplexan) | ETH2_0718500^c^ | tilS, PP-loop family protein |
| *Babesia microti* (Apicomplexan) | BMR1_01G01110^d^ | tilS, PP-loop family protein |
| *Theileria annulata (*Apicomplexan) | TA03600^d^ | tilS, hypothetical protein |

**Supplementary Table 3.** **tRNAs used for phylogenetic analysis presented in Supplementary Figure 5.** Accession ID for apicoplast genome-encoded tRNAs are provided. *Pfal*, *Plasmodium falciparum*; *Tgon*, *Toxoplasma gondii*; *Eten*, *Eimeria tenella*; *Bmic*, *Babesia microti*; *Tpar*, *Theileria parva*; *Hmar*, *Haloarcula* *marismortui*; *Mmob*, *Mycoplasma mobile*; *Bsub*, *Bacillus subtilis*; *Ecol*, *Escherichia coli*; *Cpan*, *Cycas panzhihuaensis*; *Slyc*, *Solanum lycopersicum*; *Scer*, *Saccharomyces* *cerevisiae*.

| **tRNA name** | **Accession ID** | **Sequence (5' → 3')** |
| --- | --- | --- |
| *Pfal*_trnM-CAU1 | PF3D7_API06600 | AGCGAAAUAGAGCAUAAGGAAAGUUCGUC  GGAUUCAUGCUCCGAAGGUAAUCGGUUC  AAUUCCGUUUUUCGCUUA |
| *Pfal*_trnM-CAU2 | PF3D7_API00600 | AACAUUUAUAGCUAAGUGGUCGAAAGCAA  UGGACUCAUAAUUCAUUUUCAUAUAUUGA  UCAUCAGUAGUUCGAAUCUACUUAAAUGU |
| *Pfal*_trnM-CAU3 | PF3D7_API05000 | AGCGAAAUAGAGCAUAAGGAAAGUUCGUC  GGAUUCAUGCUCCGAAGGUAAUCGGUUC  AAUUCCGUUUUUCGCUU |
| *Pfal*_trnI-GAU | PF3D7_API05800 | AUAGGUUUUUAGUUUAAUGGUUAAAACAU  ACUCUUGAUAAGGGUAAAAUUUUAGUUCA  AUUCUAAAAUAACC |
| *Tgon*_trnM-CAU1 | TGME49_355180 | AGCGGGGUAGAGCAGGUUGGUAGCUCGU  CGGGCUCAUGACCCGAAGGUCAGCGGUU  CAAAUCCGCUCCUCGUUU |
| *Tgon*_trnM-CAU2 | TGME49_355060 | AUACUUGUGGCUGAGUGGGCAAAAGCAGU  GAGCUCAUAACUCAUAUAAAACGAAAGUUC  GAAUCUUUUCAAGUAUA |
| *Tgon*_trnI-GAU1 | TGME49_355100 | AGGCUAGUAGCUCAACGGUAGAGCACGCU  UUUGAUAAGGGCGUGGUUUCUGGUUCGA  UUCCAGGGUGGCCUA |
| *Tgon*_trnI-GAU2 | TGME49_355110 | AGGCUAGUAGCUCAACGGUAGAGCACGCU  UUUGAUAAGGGCGUGGUUUCUGGUUCGA  UUCCAGGGUGGCCUA |
| *Eten*_trnM-CAU1 | ETH2_API00800 | AAACGGAGUAGAGCAGUCUGGUUAGC  UCAUCGGGCUCAUGAUCCGAAGGUCA  ACGGUUCAAUUCCGUUCUCCGUUUU |
| *Eten*_trnM-CAU2 | ETH2_API01500 | UGUACCUGUGGCUGAGUGGUCAAAAG  CGGUGGGCUCAUAAUCCAUUUUUUUU  CAAAAGUUCAAAUCUUUUCAGGUAUAA |
| *Eten*_trnM-CAU3 | ETH2_API05600 | AAACGGAGUAGAGCAGUCUGGUUAGCU  CAUCGGGCUCAUGAUCCGAAGGUCAAC  GGUUCAAUUCCGUUCUCCGUUUU |
| *Eten*_trnI-GAU1 | ETH2_API00100 | GGGCUGUUAGCUCAUCGGUAGAGCGCG  CCCCUGAUAAGGGCGAGGUACCUGGUU  CAACCCCAGGACGGCCUA |
| *Eten*_trnI-GAU2 | ETH2_API06300 | GGGCUGUUAGCUCAUCGGUAGAGCGCG  CCCCUGAUAAGGGCGAGGUACCUGGUU  CAACCCCAGGACGGCCUA |
| *Bmic*_trnM-CAU1 | BmR1_api00060 | AAUAAGAUAUAGUAAUAAGGAAACUUACC  AGCUUCAUGGUCUGGAGAUUGCAGUUC  GAGUCUGCAUCUUAUUU |
| *Bmic*_trnM-CAU2 | BmR1_api00070 | AUAUCUGUAGCUUAGUGGUUAUAGCAAU  GGGCCCAUGACUCAUUAAUUUCAGUAGU  UCAAAUCUACUCAGAUAUA |
| *Bmic*_trnM-UAU | BmR1_api00200 | AAUAAUUUAUUAUUUUAUAUAUAUAUAUU  AUAUAUAUAUAAAAUAAUAAAUAUUU |
| *Bmic*_trnI-GAU | BmR1_api00090 | AGAUUUUUAGUUUACUGGUAAAACAUAUC  UUUGAUAAGGAUAAAAUAUUUGGUUCAAU  UCCAAAAAAAUCUA |
| *Tpar*_trnM-CAU | TpMuguga_05g00051 | GUAUCUAUAGCUUAGAGGCUAAAGCGAU  GAGUUCAUACCUCAUGUACAGUAGUUCA  AAUCUACUUAGAUAUA |
| *Tpar*_trnI-GAU | TpMuguga_05g00069 | GGACUUUUAGCUUAAUUGUUAAAGUUUA  CAUGUGAUAUAUGUGAGAGUUUUGGUUA  AAAUCCAAAAAAGUCCA |
| *Hmar*_trnM-CAU |  | GCCCGGGUGGCUUAGCUGGACAUAGCG  CCGCACUCAUAAUGCGGAGAUCGAGGG  UUCGGA |
| *Hmar*_trnfM-CAU |  | AGCGGGAUGGGAUAGCCAGGAGAUUCCG  GCGGGCUCAUAACCCGCAGAUCGGUAGU  UCAAAUCUACCUCCCGCUA |
| *Hmar*_trnI-CAU |  | GGGCCCUUAGCUUAGUCUGGUUAAAGCG  AUCGGCUCAUAACCGAUUGAGCGCUGGU  UCAAAUCCGGCAGGGCCCA |
| *Hmar*_trnI-GAU |  | GGGCCAAUAGCUCAAUCAGGUUGAGCGC  UCGGCUGAUAACCGGGAGGUUCGCGGUU  CAAAUCCGCGUUGGCCCA |
| *Mmob*_trnM-CAU |  | GGCUCUGUAGCUCAGCUGGUUAGAGCAU  UCGGUUCAUACCCGAAAGGUCAAGAGUUC  GACUCUCUUCGGAGCUACCA |
| *Mmob*_trnI-GAU |  | GGGAGCGUAGCUCAGCUGGUUAGAGCAC  ACGACUGAUAAUCGUGAGGUCGAUGGUU  CGAGUCCAUUCGUUCCCACCA |
| *Mmob*_trnI-UAU |  | GGUCCUAUAGCUCAGUCGGUUAGAGCAC  ACGACUUAUAAUCGUGAGGUCGCUGGUU  CAAUCCCAGCUAGGACUACCA |
| *Bsub*_trnM-CAU1 |  | GGCGGUGUAGCUCAGCUGGCUAGAGCGU  ACGGUUCAUACCCGUGAGGUCGGGGGUU  CGAUCCCCUCCGCCGCUACCA |
| *Bsub*_trnM-CAU2 |  | GGCGGUGUAGCUCAGCUGGCUAGAGCG  UACGGUUCAUACCCGUGAGGUCGGGGG  UUCGAUCCCCUCCGCCGCUACCA |
| *Bsub*_trnfM-CAU1 |  | CGCGGGGUGGAGCAGUUCGGUAGCUCG  UCGGGCUCAUAACCCGAAGGUCGCAGGU  UCAAAUCCUGCCCCCGCAACCA |
| *Bsub*_trnfM-CAU2 |  | CGCGGGGUGGAGCAGUUCGGUAGCUCG  UCGGGCUCAUAACCCGAAGGUCGCAGGU  UCAAAUCCUGCCCCCGCAACCA |
| *Bsub*_trnfM-CAU3 |  | CGCGGGGUGGAGCAGUUCGGUAGCUCGU  CGGGCUCAUAACCCGAAGGUCGCAGGUUC  AAAUCCUGCCCCCGCAACCA |
| *Bsub*_trnI2-CAU1 |  | GGACCUUUAGCUCAGUUGGUUAGAGCAGA  CGGCUCAUAACCGUCCGGUCGUAGGUUCG  AGUCCUACAAGGUCCACCA |
| *Bsub*_trnI-GAU1 |  | GGGCCUGUAGCUCAGCUGGUUAGAGCGCA  CGCCUGAUAAGCGUGAGGUCGAUGGUUCG  AGUCCAUUCAGGCCCACCA |
| *Bsub*_trnI2-GAU1 |  | GGGCCUGUAGCUCAGCUGGUUAGAGCGCA  CGCCUGAUAAGCGUGAGGUCGGUGGUUCG  AGUCCACUCAGGCCCACCA |
| *Bsub*_trnI2-GAU2 |  | GGGCCUGUAGCUCAGCUGGUUAGAGCGCAC  GCCUGAUAAGCGUGAGGUCGGUGGUUCGAG  UCCACUCAGGCCCACCA |
| *Ecol*_trnfM-CAU1 |  | CGCGGGGUGGAGCAGCCUGGUAGCUC  GUCGGGCUCAUAACCCGAAGGUCGUCG  GUUCAAAUCCGGCCCCCGCAACCA |
| *Ecol*_trnfM-CAU2 |  | CGCGGGGUGGAGCAGCCUGGUAGCUC  GUCGGGCUCAUAACCCGAAGGUCGUC  GGUUCAAAUCCGGCCCCCGCAACCA |
| *Ecol*_trnfM-CAU3 |  | CGCGGGGUGGAGCAGCCUGGUAGCUC  GUCGGGCUCAUAACCCGAAGGUCGUCG  GUUCAAAUCCGGCCCCCGCAACCAA |
| *Ecol*_trnfM-CAU4 |  | CGCGGGGUGGAGCAGCCUGGUAGCUCG  UCGGGCUCAUAACCCGAAGAUCGUCGGU  UCAAAUCCGGCCCCCGCAACCA |
| *Ecol*_trnM-CAU1 |  | GGCUACGUAGCUCAGUUGGUUAGAGCAC  AUCACUCAUAAUGAUGGGGUCACAGGUU  CGAAUCCCGUCGUAGCCACCA |
| *Ecol*_trnM-CAU2 |  | GGCUACGUAGCUCAGUUGGUUAGAGCACA  UCACUCAUAAUGAUGGGGUCACAGGUUCG  AAUCCCGUCGUAGCCACCA |
| *Ecol*_trnI-GAU1 |  | AGGCUUGUAGCUCAGGUGGUUAGAGCGCA  CCCCUGAUAAGGGUGAGGUCGGUGGUUCA  AGUCCACUCAGGCCUACCA |
| *Ecol*_trnI-GAU2 |  | AGGCUUGUAGCUCAGGUGGUUAGAGCGCA  CCCCUGAUAAGGGUGAGGUCGGUGGUUC  AAGUCCACUCAGGCCUACCA |
| *Ecol*_trnI2-CAU1 |  | GGCCCCUUAGCUCAGUGGUUAGAGCAGG  CGACUCAUAAUCGCUUGGUCGCUGGUUC  AAGUCCAGCAGGGGCCACCA |
| *Ecol*_trnI2-CAU2 |  | GGCCCUUUAGCUCAGUGGUUAGAGCAGG  CGACUCAUAAUCGCUUGGUCGCUGGUUC  AAGUCCAGCAAGGGCCACCA |
| *Ecol*_trnI-GAU3 |  | AGGCUUGUAGCUCAGGUGGUUAGAGCGC  ACCCCUGAUAAGGGUGAGGUCGGUGGUU  CAAGUCCACUCAGGCCUACCA |
| *Cpan*_trnM-CAU1 |  | ACCUACUUAACUCAGUGGUUAGAGUAUCG  CUUUCAUACGGCGGGAGUCAUUGGUUCA  AAUCCAAUAGUAGGUA |
| *Cpan*_trnM-CAU2 |  | UGCGGGGUAGAGCAGUUUGGUAGCUCGC  AAGGCUCAUAACCUUGAGGUCACGGGUUC  AAAUCCCGUCUCCGCCA |
| *Cpan*_trnI-GAU1 |  | UGGGCUAUCCUGGACUUGAACCAGAGACC  UCGCCCGUAUCAGGGGCGCGCUCUACCA  CUGAGCUAAUAGCCC |
| *Cpan*_trnI-GAU2 |  | GGGCUAUUAGCUCAGUGGUAGAGCGCGCC  CCUGAUGGGCGAGGUCUCUGGUUCAAGUC  CAGGAUAGCCCA |
| *Cpan*_trnI-AUA |  | GCAUCCAUGGCUGAACGGUUAAAGCGCCC  AACUCAUAAUUGGCGAAUUCGCAGGUUCAA  UUCCUGCUGGAUGCA |
| *Slyc*_trnfM-CAU |  | CGCGGGGUAGAGCAGUUUGGUAGCUCGCA  AGGCUCAUAACCUUGAGGUCACGGGUUCA  AAUCCUGUCUCCGCAA |
| *Slyc*_trnM-CAU |  | ACCUACUUAACUCAGUGGUUAGAGUACUG  CUUUCAUACGGCGGGAGUCAUUGGUUCAA  AUCCAAUAGUAGGUA |
| *Slyc*_trnI-CAU |  | GCAUCCAUGGCUGAAUGGUUAAAGCGCCC  AACUCAUAAUUGGCGAAUUCGUAGGUUCAA  UUCCUACUGGAUGCA |
| *Slyc*_trnI-GAU |  | GGGCUAUUAGCUCAGUGGUAGAGCGCGCC  CCUGAUAAUUGCGGGGCGAGGUCUCUGGU  UCAAGUCCAGGAUGGCCCA |
| *Scer*_trniM-CAU1 |  | AGCGCCGUGGCGCAGUGGAAGCGCGCAGG  GCUCAUAACCCUGAUGUCCUCGGAUCGAAA  CCGAGCGGCGCUA |
| *Scer*_trniM-CAU2 |  | AGCGCCGUGGCGCAGUGGAAGCGCGCAGGG  CUCAUAACCCUGAUGUCCUCGGAUCGAAACC  GAGCGGCGCUA |
| *Scer*_trniM-CAU3 |  | AGCGCCGUGGCGCAGUGGAAGCGCGCAGGG  CUCAUAACCCUGAUGUCCUCGGAUCGAAACC  GAGCGGCGCUA |
| *Scer*_trniM-CAU4 |  | AGCGCCGUGGCGCAGUGGAAGCGCGCAGGG  CUCAUAACCCUGAUGUCCUCGGAUCGAAACC  GAGCGGCGCUA |
| *Scer*_trniM-CAU5 |  | AGCGCCGUGGCGCAGUGGAAGCGCGCAGGG  CUCAUAACCCUGAUGUCCUCGGAUCGAAACC  GAGCGGCGCUA |
| *Scer*_trnM-CAU1 |  | GCUUCAGUAGCUCAGUAGGAAGAGCGUCAGU  CUCAUAAUCUGAAGGUCGAGAGUUCGAACCU  CCCCUGGAGCA |
| *Scer*_trnM-CAU2 |  | GCUUCAGUAGCUCAGUAGGAAGAGCGUCAG  UCUCAUAAUCUGAAGGUCGAGAGUUCGAAC  CUCCCCUGGAGCA |
| *Scer*_trnM-CAU3 |  | GCUUCAGUAGCUCAGUAGGAAGAGCGUCAG  UCUCAUAAUCUGAAGGUCGAGAGUUCGAAC  CUCCCCUGGAGCA |
| *Scer*_trnM-CAU4 |  | GCUUCAGUAGCUCAGUAGGAAGAGCGUCAG  UCUCAUAAUCUGAAGGUCGAGAGUUCGAAC  CUCCCCUGGAGCA |
| *Scer*_trnM-CAU5 |  | GCUUCAGUAGCUCAGUAGGAAGAGCGUCAG  UCUCAUAAUCUGAAGGUCGAGAGUUCGAAC  CUCCCCUGGAGCA |
| *Scer*_trnM-CAU6 |  | UGCAAUAUGAUGUAAUUGGUUAACAUUUUAG  GGUCAUGACCUAAUUAUAUACGUUCAAAUCG  UAUUAUUGCUA |
| *Scer*_trnM-CAU7 |  | GCUUGUAUAGUUUAAUUGGUUAAAACAUUUG  UCUCAUAAAUAAAUAAUGUAAGGUUCAAUUCC  UUCUACAAGUA |
| *Scer*_trnI-AAU1 |  | GGUCUCUUGGCCCAGUUGGUUAAGGCACCG  UGCUAAUAACGCGGGGAUCAGCGGUUCGAUC  CCGCUAGAGACCA |
| *Scer*_trnI-AAU2 |  | GGUCUCUUGGCCCAGUUGGUUAAGGCACCG  UGCUAAUAACGCGGGGAUCAGCGGUUCGAU  CCCGCUAGAGACCA |
| *Scer*_trnI-AAU4 |  | GGUCUCUUGGCCCAGUUGGUUAAGGCACCG  UGCUAAUAACGCGGGGAUCAGCGGUUCGAU  CCCGCUAGAGACCA |
| *Scer*_trnI-AAU3 |  | GGUCUCUUGGCCCAGUUGGUUAAGGCACC  GUGCUAAUAACGCGGGGAUCAGCGGUUCG  AUCCCGCUAGAGACCA |
| *Scer*_trnI-AAU5 |  | GGUCUCUUGGCCCAGUUGGUUAAGGCACC  GUGCUAAUAACGCGGGGAUCAGCGGUUCG  AUCCCGCUAGAGACCA |
| *Scer*_trnI-AAU6 |  | GGUCUCUUGGCCCAGUUGGUUAAGGCACC  GUGCUAAUAACGCGGGGAUCAGCGGUUCGA  UCCCGCUAGAGACCA |
| *Scer*_trnI-AAU7 |  | GGUCUCUUGGCCCAGUUGGUUAAGGCACC  GUGCUAAUAACGCGGGGAUCAGCGGUUCGA  UCCCGCUAGAGACCA |
| *Scer*_trnI-AAU9 |  | GGUCUCUUGGCCCAGUUGGUUAAGGCACC  GUGCUAAUAACGCGGGGAUCAGCGGUUCGA  UCCCGCUAGAGACCA |
| *Scer*_trnI-AAU8 |  | GGUCUCUUGGCCCAGUUGGUUAAGGCACC  GUGCUAAUAACGCGGGGAUCAGCGGUUCG  AUCCCGCUAGAGACCA |
| *Scer*_trnI-AAU10 |  | GGUCUCUUGGCCCAGUUGGUUAAGGCACC  GUGCUAAUAACGCGGGGAUCAGCGGUUCGA  UCCCGCUAGAGACCA |
| *Scer*_trnI-AAU11 |  | GGUCUCUUGGCCCAGUUGGUUAAGGCACC  GUGCUAAUAACGCGGGGAUCAGCGGUUCGA  UCCCGCUAGAGACCA |
| *Scer*_trnI-AAU12 |  | GGUCUCUUGGCCCAGUUGGUUAAGGCACC  GUGCUAAUAACGCGGGGAUCAGCGGUUCG  AUCCCGCUAGAGACCA |
| *Scer*_trnI-AAU13 |  | GGUCUCUUGGCCCAGUUGGUUAAGGCACC  GUGCUAAUAACGCGGGGAUCAGCGGUUCG  AUCCCGCUAGAGACCA |
| *Scer*_trnI-GAU |  | GAAACUAUAAUUCAAUUGGUUAGAAUAGUAU  UUUGAUAAGGUACAAAUAUAGGUUCAAUCCC  UGUUAGUUUCA |
| *Scer*_trnI-UAU1 |  | GCUCGUGUAGCUCAGUGGUUAGAGCUUCGU  GCUUAUAGCAACAUUCGGUUUCCGAAGUUUC  UGUGCCAAAGACCUUUCAAACAGGCCUUUAA  AAGCAACGCGACCGUCGUGGGUUCAAACCCC  ACCUCGAGCA |
| *Scer*_trnI-UAU2 |  | GCUCGUGUAGCUCAGUGGUUAGAGCUUCGU  GCUUAUAGCAACAUUCGGUUUCCGAAGUUUC  UGUGCCAAAGACCUUUCAAACAGGCCUUUAA  AAGCAACGCGACCGUCGUGGGUUCAAUCCCC  ACCUCGAGCA |
